# Pan-genome analysis of *Mycobacterium tuberculosis* identifies accessory genome sequences deleted in modern Beijing lineage

**DOI:** 10.1101/2020.12.01.407569

**Authors:** Syed Beenish Rufai, Egon A. Ozer, Sarman Singh

## Abstract

Beijing sub-lineage of *Mycobacterium tuberculosis* has been reported to have increased transmissibility and drug resistance. This led us to get insights of genomic landscape of modern Beijing sub-lineages in comparison with other lineages of *M. tuberculosis* utilizing pan-genomics approach. Pangenome analysis was performed using software Spine (v0.2.3), AGEnt (v0.2.3) and ClustAGE (v0.7.6). The average pangenome size was 45,40,849 bp with 4,391 coding sequences (CDS), with a GC content of 65.4%. The size of the core genome was 36,83,161 bp, contained 3,698 CDS and had an average GC content of 65.1%. The average accessory genome size was 6,96,320.9 bp, with 539.4 CDS and GC content of 67.9%. Among the accessory elements complete deletion of CRISPR-associated endoribonuclease cas1 (*Rv2817c*), cas2 (*Rv2816c*), CRISPR type III-a/mtube-associated protein csm6 (*Rv2818c*), CRISPR type III-a/mtube-associated ramp protein csm5 (*Rv2819c*) and partial deletion (61.5%) CRISPR type III-a/mtube-associated ramp protein csm4 (*Rv2820c*) sequences was found specifically in modern Beijing lineages taken in assortment. The sequences were validated using conventional PCR method, which precisely amplified the corresponding targets of sequence elements with 100% sensitivity and specificity. Deletion of accessory CRISPR sequence elements amongst the modern Beijing sub-lineage of *M. tuberculosis* suggest more defective DNA-repair in these strains which may enhance virulence of the strains. Further, the developed conventional PCR approach for detection of virulent modern Beijing lineage may be of interest to public health and outbreak control organizations for rapid detection of modern Beijing lineage.

## Introduction

The emergence of resistance towards first and second line anti-tuberculosis drugs in *M. tuberculosis* strains poses an increasing threat to public health (1). Estimates from TB investigation data predicts an expected increase in Multi-drug resistant (MDR) and extensively drug resistant (XDR) TB especially in developing countries like India, Philippines, Russia and South Africa (2). Classical techniques of determining genotyping and recently Whole genome sequencing (WGS) have revealed significant evolution of strains leading to diversity among human adapted *M. tuberculosis* strains, leading to acquire various genetic mechanisms for successful transmission and survival in the population (3).

Among different lineage of lineages of *M. tuberculosis* which include lineages Indo Oceanic, Euro-American, Central Asian, and the East Asian (Beijing-sub lineage) is known to have originated out of East Asia and has disseminated around the world. The variability in gene expression patterns of different strains of *M. tuberculosis* during infection and intra/inter genomic variation among pathogenic strains has been documented as a significant feature in pathogenesis and types of infection caused by the strains (4). The Beijing sub-lineage is reported as more virulent in comparison to other lineages of *M. tuberculosis* in terms of an enhanced level of pathogenicity leading to increased transmissibility, rapid progression from latent to active disease, epidemiological association with transmission outbreaks and increased frequency of antibiotic resistance suggesting that genetic modifications distinguish the virulence and pathogenicity of this lineage (5–11). The global spread and association of Beijing strains with drug resistance has also been documented from India and other parts of world which is driving attention of researchers to further understand the genomic features that make this lineage more virulent in comparison to that of other lineages of *M. tuberculosis* (6, 8, 12–14).

With the advent of next-generation sequencing (NGS) technologies, WGS has been used widely to identify evolutionary markers, polymorphism across lineages and mutations associated with drug resistance in *M. tuberculosis* (15). With availability of genomic databases and whole genome sequences of *M. tuberculosis*, approaches for deciphering novel mutations in comparison to reference strains are being utilized which results in loss of unique coding sequences(CDS)/genes that may have role in virulence or acquiring drug resistance (16).To overcome this, pan-genome based approach has been preferred which discern a more complete gene landscape for identification of unique (specific to single strains), accessory (shared among two or more strains) and core genes (present in all strains) for estimating the genomic diversity and identification of novel/unique gene sequences and discovery of markers for lineage identification (16–18).

WGS of *M. tuberculosis* isolates has been performed on isolates from Malaysia, China, Myanmar, Peru, Colombia, India, Ireland, New Zealand, Africa, Korea and Russia and large collection of genomes from clinical isolates of *M. tuberculosis* were analyzed by Manson et al 2017 [21–25]. But no pangenome studies were done on *M. tuberculosis* diversity of lineages to understand the variations in terms of unique genes/ sequences among them. This study was aimed to identify unique and shared sequences in the three laboratory sequenced TB isolates and WGS *M. tuberculosis* genomes data available in the public domain to decipher novel markers that may be questioned for higher virulence in the Beijing lineage.

## Material and Methods

This study was performed in an accredited TB laboratory in the Division of Clinical Microbiology & Molecular Medicine, All India Institute of Medical Sciences, New Delhi. Representative XDR-TB (L-823, L-182 and L-31) isolates of known drug susceptibility testing (DST) patterns and spoligotypes published previously were selected for WGS (19).

### Whole genome sequencing of three lab isolated XDR-TB

Whole genome sequencing of the three XDR-TB isolates was performed using Ion Torrent PGM platform (Life Technologies). Briefly, DNA extraction was performed as previously described (20). The three genomes were sequenced using Ion Torrent PGM as published previously (19). Sequencing data generated by Ion Torrent PGM was analyzed using the Torrent Suite Software. De-novo assembly of sequenced data was performed using ion-plugin-assembler St. Petersburg genome assembler (SPades) (v 3.1.0) (19, 21). The three whole genomes sequences were deposited in GenBank under the accession numbers NDYV00000000 for L-182 (beijing strain), NCTW00000000 for L-823 (beijing strain) and NDYU00000000 L-31 (central Asian strain).

### Data mining of whole Genome Draft sequences of *M. tuberculosis* isolates from the NCBI genome database

In order to understand complete genomic repertoire, analysis of different *M. tuberculosis* genomes was required for in depth analysis. Due to lack of funding we only did WGS of representative XDR-TB isolates. We explored the work of *Manson et al., 2017* which already have analyzed more than five thousand genomes different lineages and variable geographical diversity (Manson et al., 2017) and selected 91 genomes based on diversity of lineages and geographical locations (22). We performed random search in (https://www.ncbi.nlm.nih.gov/sra) using the search terms “*Mycobacterium tuberculosis*” and selected 25 genomes recently published from diverse geographical locations. A total 121 genomes were taken for pan-genome analysis as per the available computational feasibility in our setup. Among 121 draft genome assemblies, three were sequenced XDR-TB genomes at our setting, 116 were draft genome assemblies downloaded from NCBI and two were reference strains of *M. tuberculosis* under accession number NC_000962.3 and AL12345.6 respectively **(Supplementary Table 1)**.

### Pan-genome analysis of 121 isolates

Pan-genome analysis of isolates was performed using the software suite Spine, AGEnt and ClustAGE (17, 23). This software identifies the nucleotide sequences and associated annotations of the core, accessory and unique genome fractions of a sequenced strain population.

Spine v 0.2.3 was used for the identification of the conserved core genome sequence of the set of 121 *M. tuberculosis* genomes using H37Rv (NC_00926.3) as the reference genome sequence for a strict core genome (genomic sequence present in 100% of the strains) and using AL12345.6 as the reference genome for extraction of soft core genome (core genome sequence present in at least 90% of the strains) (17). AGEnt v0.2.3 was used for identifying accessory genomic elements (AGEs) in bacterial genomes by using an in-silico subtractive hybridization approach against a core genome generated using the Spine algorithm (17). ClustAGE (v0.7.5) was used to compare accessory genomic elements (AGEs) between genomes. The default threshold for alignments of 85% sequence identity over at least 100 bp was used (23).

### Estimation of phylogenetic tree using kSNP 3.0

kSNP3 program was used for construction of phylogenetic parsimony tree using pan-genome SNPs from a set of genome sequences without use of reference genome and parsimony tree that is estimated as a consensus of up to 100 equally parsimonious trees. Pan-genome parsimony tree was constructed using kSNP 3.0 (24). *K-chooser* was used to determine the optimum *k-mer* size, which was set at 21.

### Visualization of trees

Trees were visualized using Interactive tree of Life (iTOL) V3 with Bootstrapped values of original and resampled tree https://itol.embl.de (25).

### Functional annotation of core and accessory genomes sequences

Functional annotation of core, accessory and unique genome sequences were transferred from orthologs of taxa group Actinobacteria (Mycobacteriacae) using EggnNOG mapper v2 (26). COG letter categories obtained were patterned for functional description in COG database (27).

### Standardization of conventional PCR for validation of sequences on clinical isolates

Primers were designed using Primer 3 (V 4.1.0) for validation of sequences/ genomic fractions specific to lineage found from pangenome data analysis (28). Designed primers were obtained from Eurofins, India. DNA was isolated from clinical isolates (50 Beijing strains and 50 non-Beijing strains) and subjected to PCR as follows: 2.5 μl of 10X buffer, 500 mM KCl] supplied with 1 ml of 50 mM MgCl2, 0·5 μl of stock 10mM dNTP, 20pmol of each primer and 1.25U of Taq DNA polymerase and 5μl of template DNA. Each PCR was started with a ‘hot Start’ for 2min at 95°C followed by denaturation (25 cycles each of 15 sec at 95°C), annealing (25 cycles each of 15 sec at 55°C) and extension (25 cycles each of 45 sec at 68°C), and a final extension for 1 cycle for 5 min at 68°C in a thermal Cycler (MJ Research, USA) and amplified products were resolved through 2% agarose gel in Tris-acetate buffer.

## Results

### Description of *M. tuberculosis* data sets used in study

Of total 121 genomes used in study, apart from two reference strains, 44 (36.3%) genomes were of African origin, 40 (31.4%) from Asia, 29 (23.1%) from Europe and 6 (4.9%) from America. Place of origin and accession numbers of all draft genomes are mentioned in **(Supplementary Table 1)**.

### Genome assembly statistics of three XDR-TB isolates

*De novo* sequence assembly was performed using SPAdes v3.1.0 and functional annotation performed using RAST (Rapid Annotation using Subsystem Technology) yielding total genome sizes of 4,201,682, 4,288,294 and 4,311,779 bp with 65, 65.4 and 65.3% GC contents, and coding sequences of 4516, 4737 and 4813 for L-823, L-182 and L-31 respectively(19). Using spoligotyping technique, L-182 and L-823 were found to be Beijing sub-lineage of ST1 and ST236, and L-31 was found to be central Asian strain of ST1120 as per the SITVIT2 database (29).

### Construction of parsimony and core genome tree

Unrooted phylogenetic tree was constructed comparing the 121 WGS genome sequences in the context of the *M. tuberculosis* lineages circulating globally **(Figure 1)**. Total of 18,025 variable single nucleotide positions were extracted from these genome sequences to construct a phylogenetic parsimony tree. Among the genomes with known lineages, three genomes from each lineage group were taken as reference. After analysis, *M. tuberculosis* genomes were dispersed among four lineages Lineage 1(8; 6.6%), Lineage 2 (42; 34.7%), Lineage 3 (8; 6.6%) and Lineage 4 (63; 52.1%) respectively **(Supplementary Table 2)**.

**Fig. 1.**
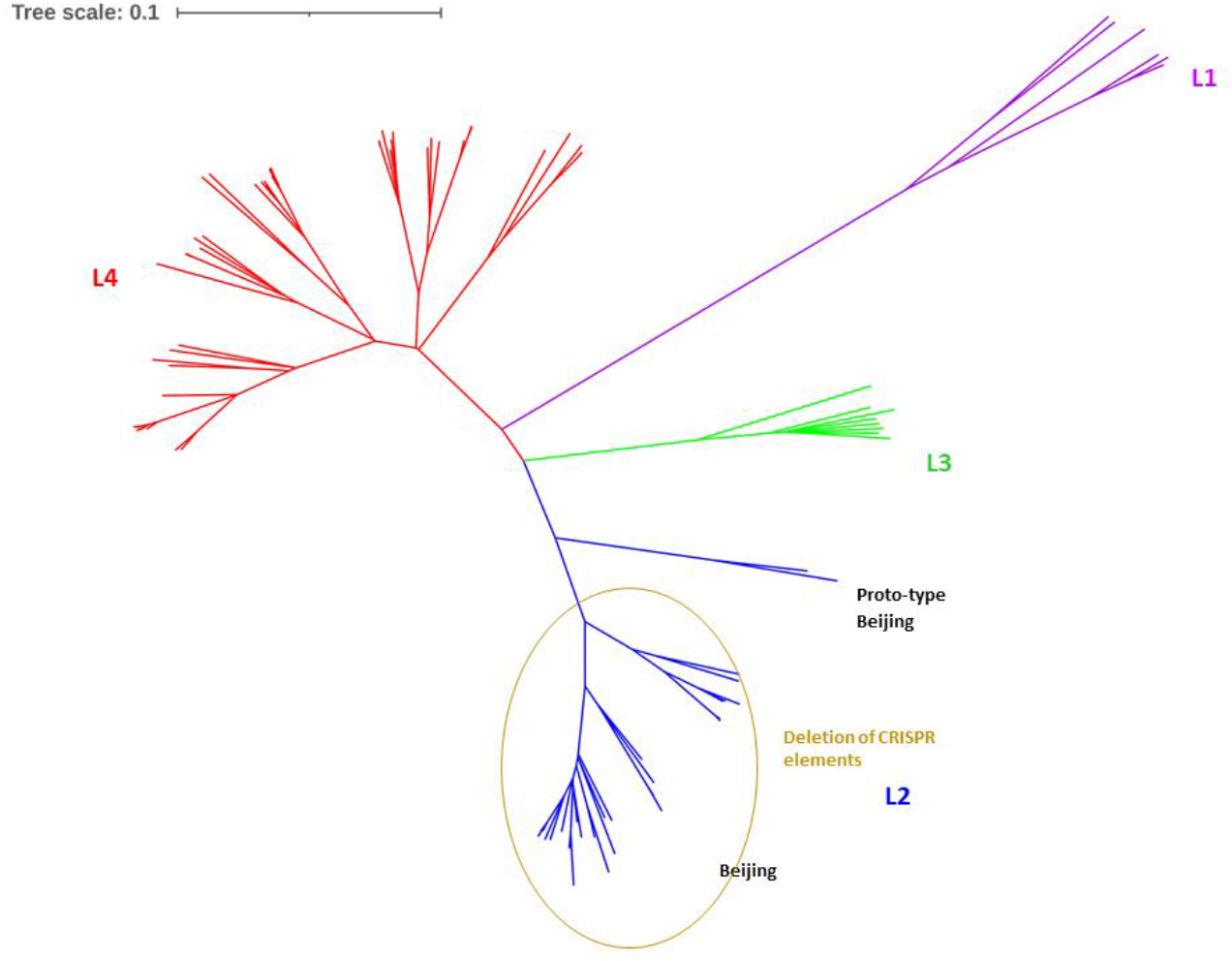
Evolutionary parsimony tree was constructed by extended Majority Rule consensus using KSNP3.0. The tree shows four distinct lineages among 121 MTB isolates. Briefly, Purple lines represent clades belonging to Lineage 1 (Indian Oceanic); Blue lines represent clades belonging to Lineage 2 [East Asian (Beijing 40; proto-type Beijing 2)]; Green lines represent clade belonging to Lineage 3 (Central Asian); Red lines represent Lineage 4 (Euro-American).

### Description of pan-genome (hard core, soft core and accessory genome)

Total pan-genome size was estimated to be 4,540,489 bp with 4391 coding sequences (CDS), and a GC content of 65.4%. Estimated average size of the hard-core genome (i.e., sequence present in 100% of genomes) was 3,683,161 bp (81.1% of total genome size), contained 3,698 coding sequences (CDS) and had an average GC content of 65.1% as compared to the average accessory genome size of 696,320.9 bp (15.3%), and GC content of 67.9%. Estimated average size of the soft-core genome (i.e. sequence present in ≥ 90% genomes) was 4,308,602 bp (94.8% of total genome size), contained 4,237 CDS and had an average GC content of 65.4% as compared to the average accessory genome size 93,819.7 bp (2.1%), with 57.6 CDS and GC content of 71.7% **(Table 1) (Supplementary Table 3)**.

**Table 1.** Estimation of hard-core, soft-core and accessory sequence elements of 121 MTB genomes.

| | Shared amongst 100% of the genome | | Shared among $\leq 90\%$ of genome | |
| --- | --- | --- | --- | --- |
| Core genome characteristics | Core genome Average (range) | Accessory genome Average (range) | Core genome Average (range) | Accessory genome Average (range) |
| Size (bp) | 3683161<br>(3672229-3751588) | 696320.9<br>(461308-742152) | 4308602<br>(4213348-44055935) | 93819.7<br>(32024-135709) |
| GC content | 65.1 | 67.9 | 65.5 | 71.7 |
| Number of sequence segments | 939<br>(894-1053) | 936.3<br>(847-1116) | 115<br>(112-658) | 121.8<br>(47-219) |
| Maximum segment length | 27136<br>(26906-33160) | 3059.4<br>(15015-27765) | 211154<br>(57005-377999) | 7391.4<br>(290.2-15599) |
| Avg. segment length | 3922.4<br>(3562.7-4170.6) | 745.6<br>(544.6-8224.4) | 37466.1<br>(6320-38375) | 778.4<br>(331.4-1026.7) |
| Median length | 2558<br>(2231-2746) | 220.2<br>(179-253) | 18237<br>(2599-20222) | 289.6<br>(71-4400) |
| No. of CDS | 3698<br>(3438-4173) | 539.3<br>(443-629) | 4237<br>(3887-4718) | 57.6<br>(20-88) |

<sup>a</sup>Core genome present in 100% of genomes
<sup>b</sup>Core genome present in $\leq 90\%$ of geno

### Determination of core genome

In order to better estimate the likely species core genome size, a rigorous definition of core genome was used. Core genome represented a total of 81.1% of the overall pan-genome repertoire. The average amount of core genome and pan-genome as a function of the number of reference genome (NC_000962.3) included in the analysis was computed using an adaptation of the method described by *Tettelin et al 2005* (16) **(Fig 2 & 3).**

**Fig 2.**
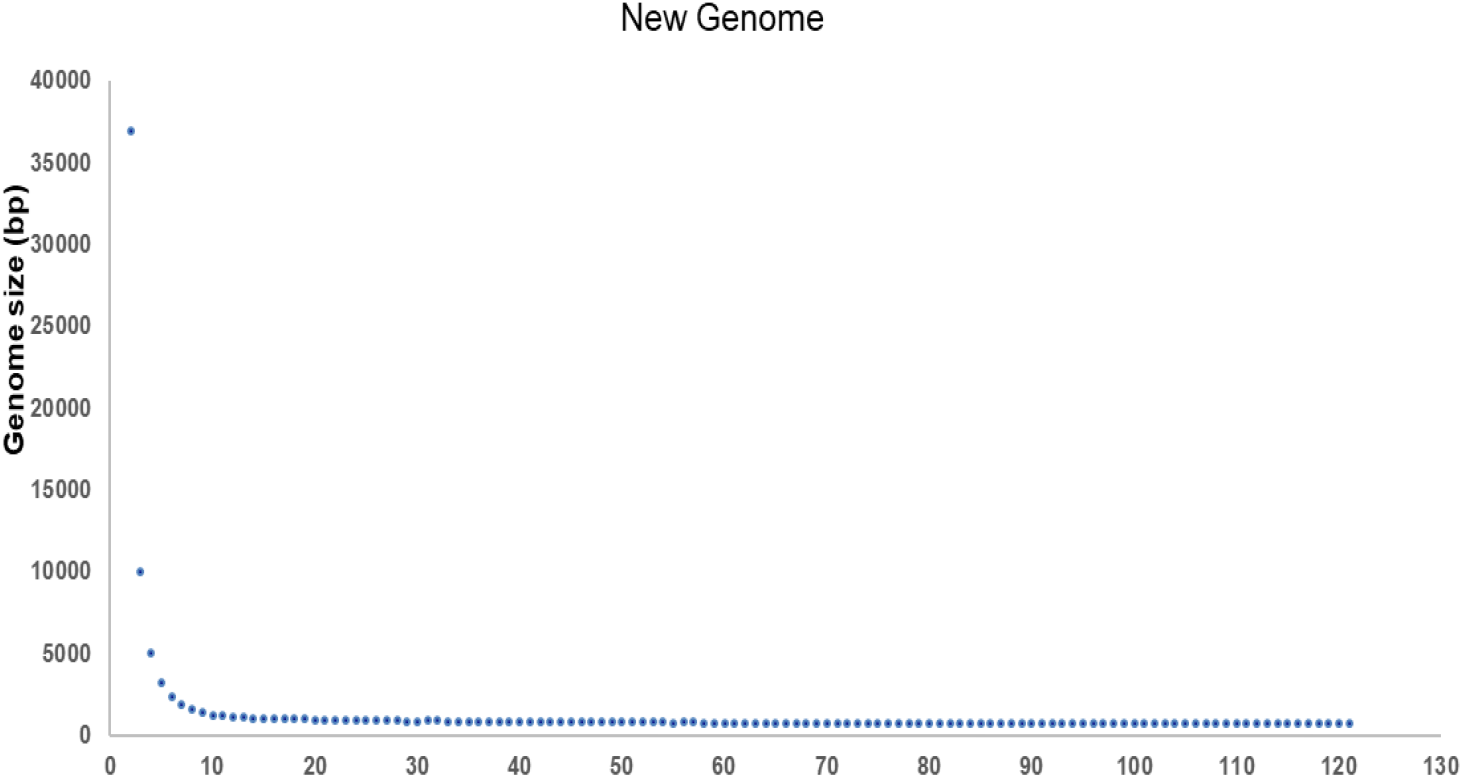
Representation of Core-Pan plot. Each marker represents the average core genome and pan-genome size of all possible permutations of genome orders for one hundred twenty for 10,000 randomly generated permutations adapted from *Tettelin et al., 2005* generated from Spine software.

**Fig. 3.**
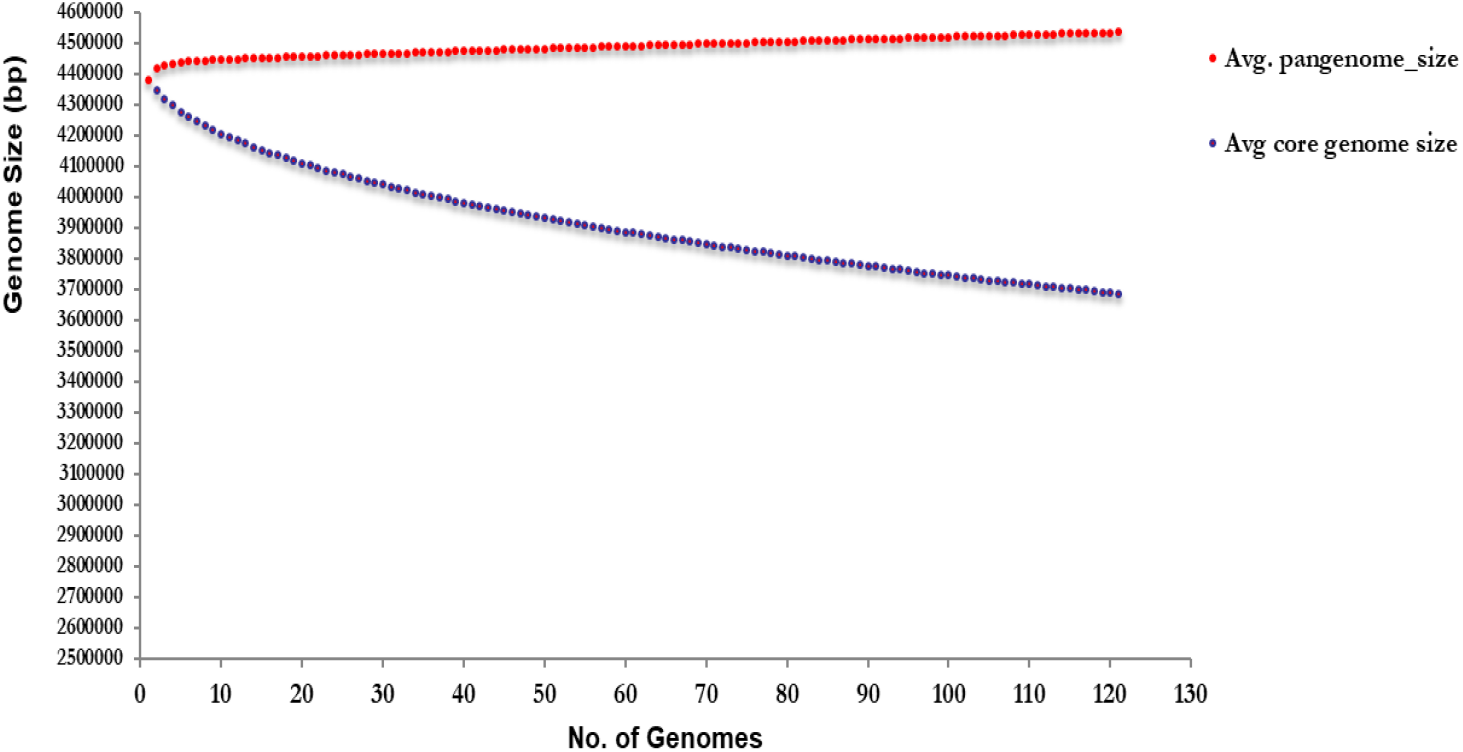
New genome size of genome orders for one hundred twenty-one for 10,000 randomly generated permutations.

### Functional annotation of core and accessory genome elements

Functional annotation of coding sequences (CDS) associated with hard-core and accessory genome were analyzed using EggNOG mapper v2. Coding sequences belonging to the core genomes were assigned to putative super-functional **(Fig 4A)** and functional **(Fig 4B)** categories using the Clusters of Orthologous Groups of proteins (COG) database. Matching of gene ontology terms with COG database predicted more than half of CDS in core genome were dedicated with metabolism functions, one fourth was of unknown function or poorly characterized and rest of CDS were involved in cellular processes and signaling, and Information storage and processing.

**Fig. 4A.**
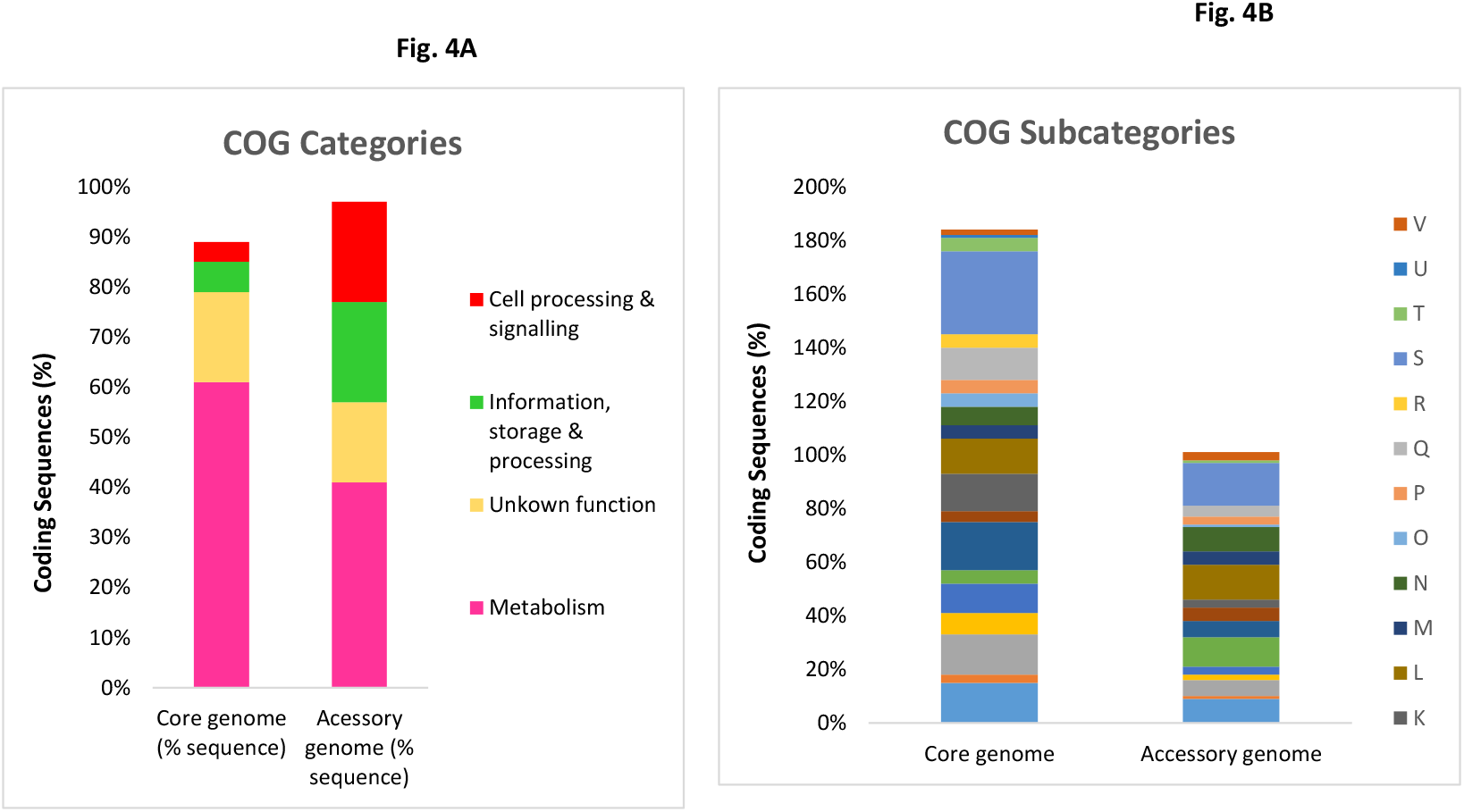
Functional annotations of core and accessory genes (A) COG categories **Fig. 4(B)** COG subcategories of predicted genes within the core and accessory genomes of *M. tuberculosis* genomes by eggNOG Mapper v2.0. Each category or subcategory is graphed as a percentage of the total number of genes in the core or accessory genomes. Sub-categories abbreviations include; 1. Cellular processes and signaling {[D] Cell cycle control, cell division, chromosome partitioning, [M] Cell wall/membrane/envelope biogenesis, [N] Cell motility, [O] Post-translational modification, protein turnover, and chaperones, [T] Signal transduction mechanisms, [U] Intracellular trafficking, secretion, and vesicular transport [V] Defense mechanisms [W] Extracellular structures [Y] Nuclear structure [Z] Cytoskeleton} 2.Information storage & processing {[A] RNA processing and modification [B] Chromatin structure and dynamics [J] Translation, ribosomal structure and biogenesis [K] Transcription [L] Replication, recombination and repair} 3.Metabolism {[C] Energy production and conversion [E] Amino acid transport and metabolism [F] Nucleotide transport and metabolism [G] Carbohydrate transport and metabolism [H] Coenzyme transport and metabolism [I] Lipid transport and metabolism [P] Inorganic ion transport and metabolism [Q] Secondary metabolites biosynthesis, transport, and catabolism} poorly characterized [R] General function prediction only [S] Function unknown

Coding sequences belonging to the accessory genomes were assigned to putative super-functional **(Fig 4A)** and functional **(Figure 4B)** categories using COG database which predicted one fourth of CDS in accessory genome were dedicated with metabolism, nearly one fourth of all CDS were associated with cellular and signaling processes, and Information storage and processing. Rest of the CDS were of unknown function.

Functional annotation of CDS associated with unique accessory elements (unique to each genome) predicted most of the genes that were acquired during the adaptation were associated with metabolism (26%), cell storage and signaling (25%), information storage and processing (20%) and rest of CDS were associated with unknown function or poorly characterized **(Fig 5)**. Variation of accessory genomic elements, among soft- and hard-core components are shown in heat map generated by ClustAGE software **(Fig. 6)**.

**Fig 5.**
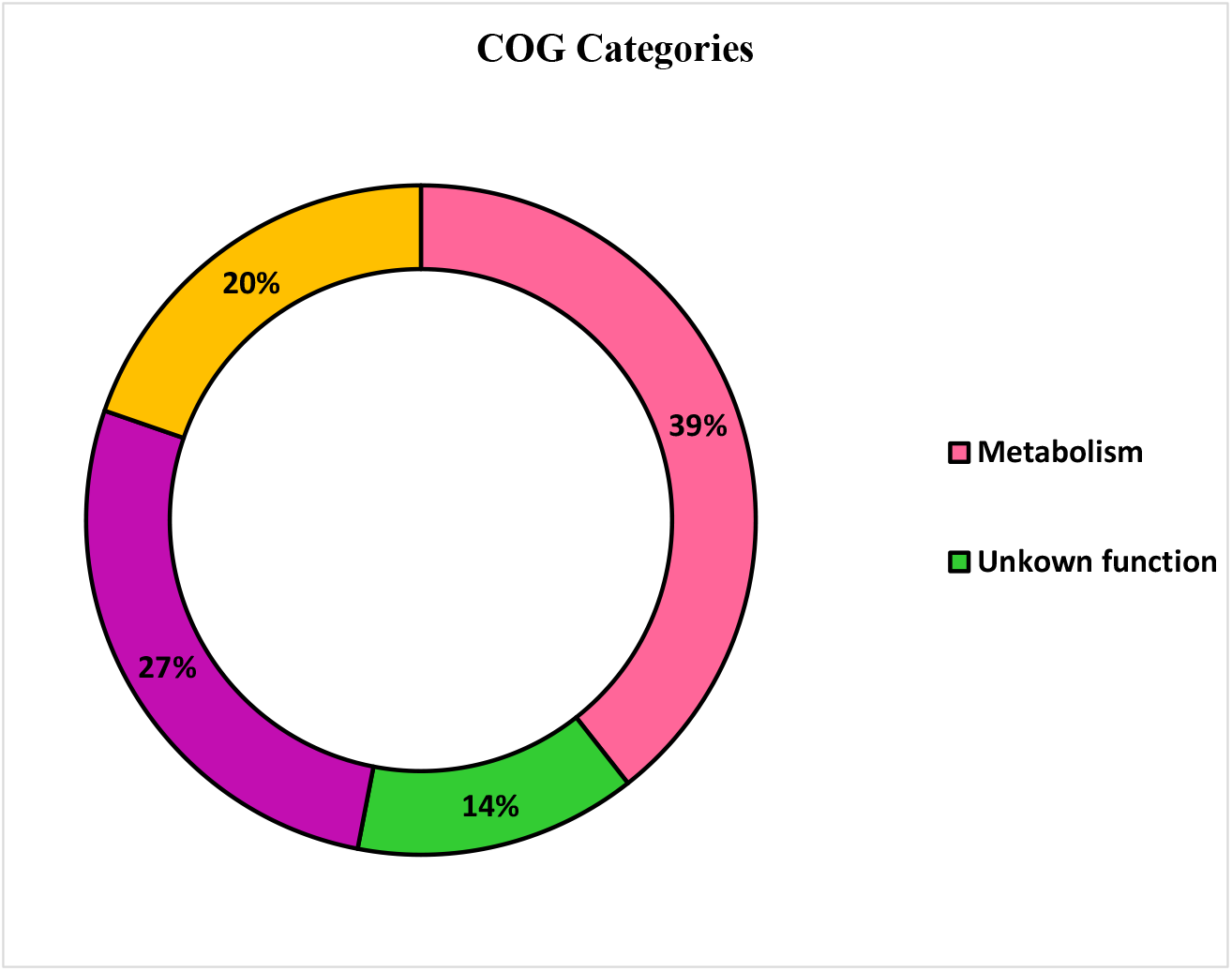
Functional annotations of unique gene COG categories of *M. tuberculosis* genomes by eggNOG Mapper v2.0.

**Fig.6.**
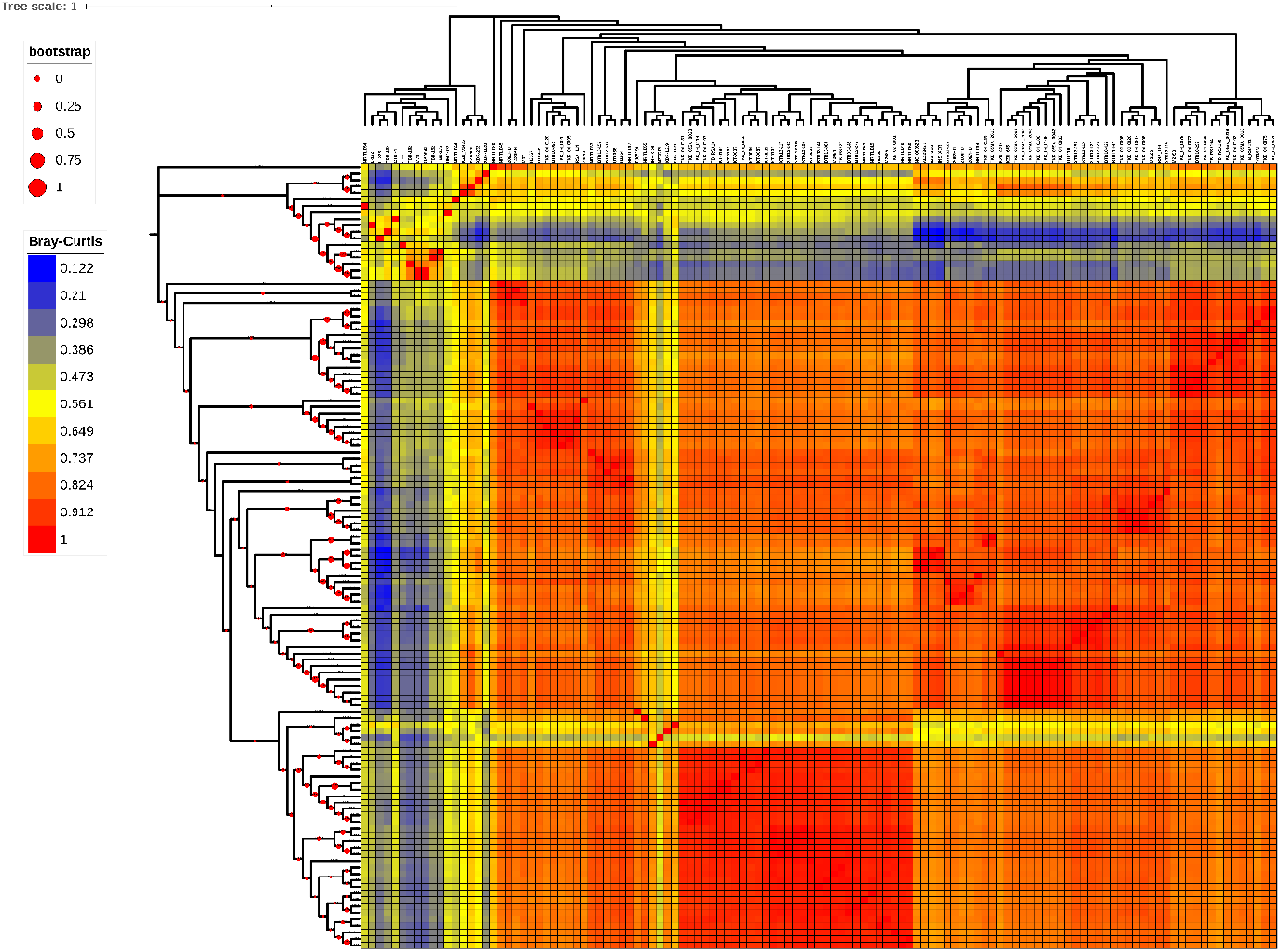
Neighbor joining tree and heat map generated by ClustAGE software showing distribution of accessory elements across 121 *M. tuberculosis* genomes. Neighbor joining tree and heat map of accessory element distribution patterns was calculated using Bray-Curtis distance matrix by ClustAGE software from distributions of accessory elements. Tree and heat map files were viewed iTOL (https://itol.embl.de)

### Clustering, distribution analysis and functional annotation of accessory nucleotide sequences

Accessory genomic elements (AGE’s) in the population, also referred to as bins, were identified using ClustAGE software (23). A total of 651 individual AGE’s hereafter referred as bins were found to be present in total input of 121 genomes that ranged in size from 201 bp to 37765 bp. The average size of the bins (651bins) was 1230.5 bp. These bins were further subdivided to sub-elements in order to see sharing and unique sequence elements among genomes. Total 859 sub-elements of sizes more than 200 bp were found in these genomes (n > 1). Shared sequences obtained from accessory elements obtained on analysing hard-core repertoire in *M. tuberculosis* strains (90%) as shown in ClustAGE plots (**Fig 7A** & **7B)**. Of the total genomes taken in assortment, 11 (9%) of *M. tuberculosis* genomes was having 139 unique sub-elements with an average length of 459.6 bp.

**Fig. 7A.**
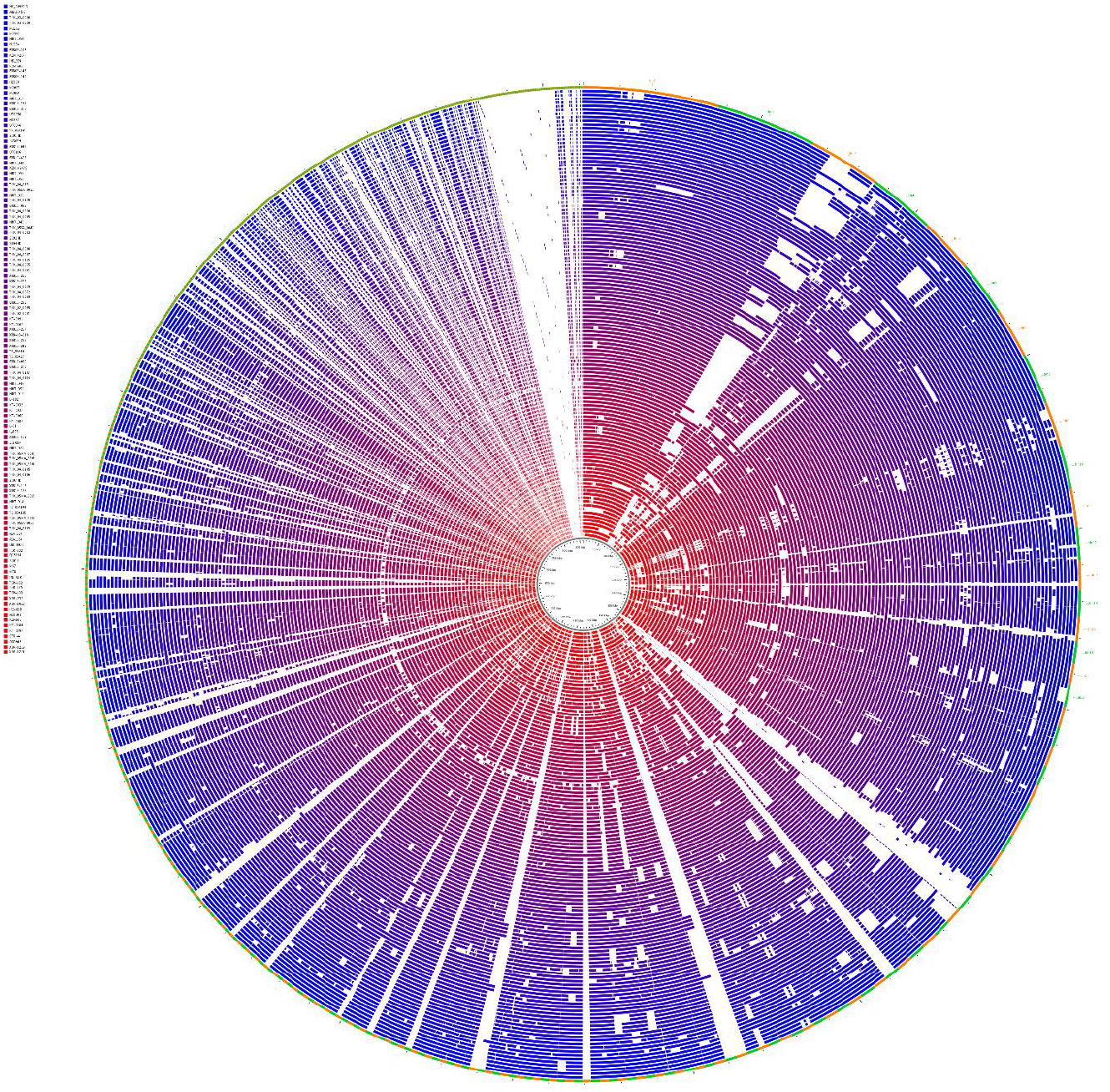
ClustAGE plot Sharing of accessory sequences present in all 121 MTB strains (100% core)

**Fig. 7B.**
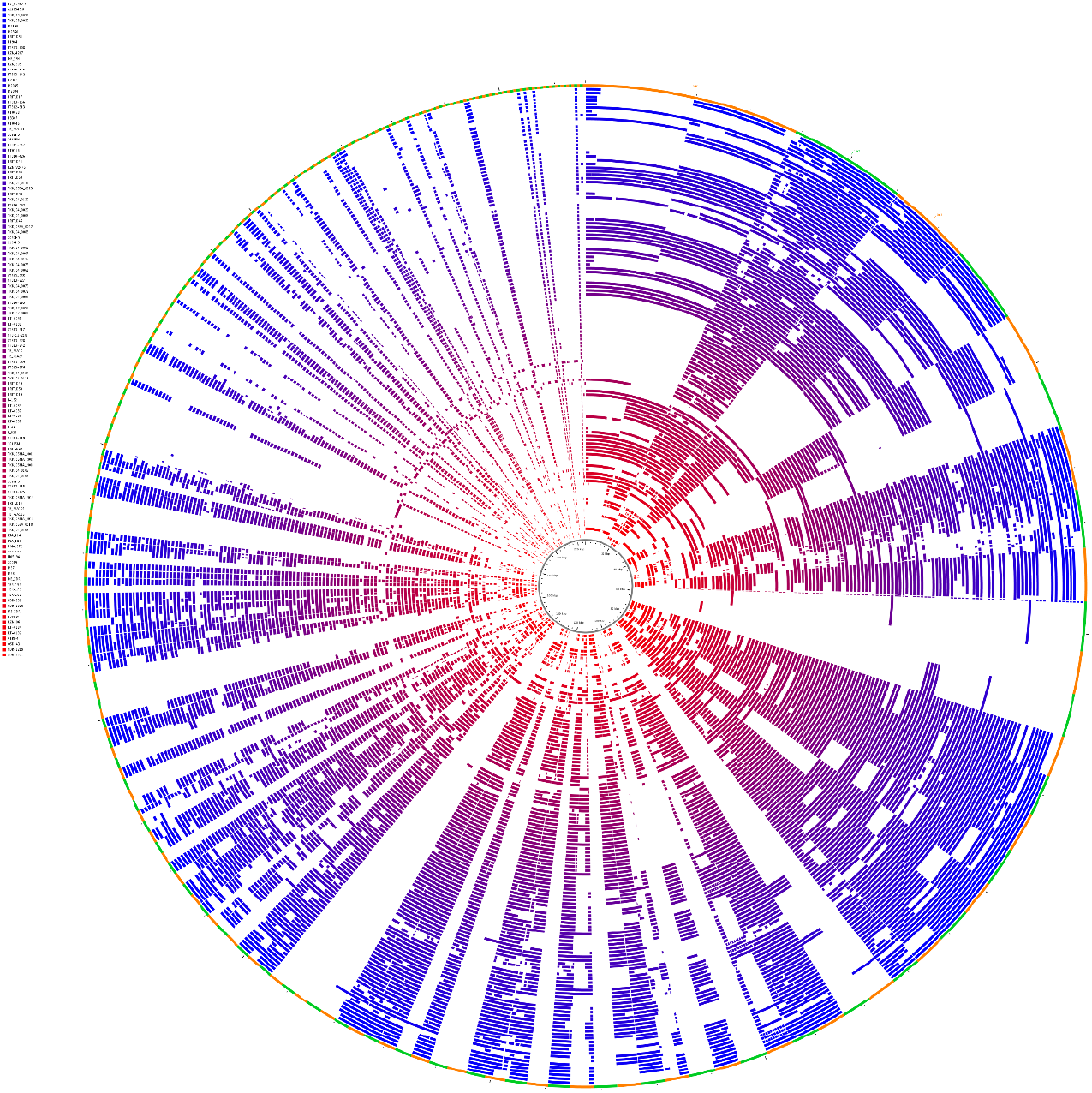
ClustAGE plot Sharing of accessory sequences of sequences or and in ≥ ≤110 MTB strains (90%

### Identification of novel markers among AGE’s

Among the shared accessory genome sequences, portions of one 4487 bp AGE (bin 32) was found to be absent in all of the Beijing genomes **(Fig 8)**. To better understand the nature of the deleted genomic sequence, bin 32 was further divided into discrete sub elements which revealed the portion of the AGE missing in the Beijing lineage strains, named bin32se-0001, encoded all or some of the following genes: CRISPR-associated endo-ribonuclease cas2 (100%); CRISPR-associated endonuclease cas1 (100%); CRISPR type III-a/mtube-associated protein csm6 (100%); CRISPR type III-a/mtube-associated protein csm5 (100%;) and CRISPR type III-a/mtube-associated ramp protein csm4 (61.5%) **(Table 2).**

**Fig 8.**
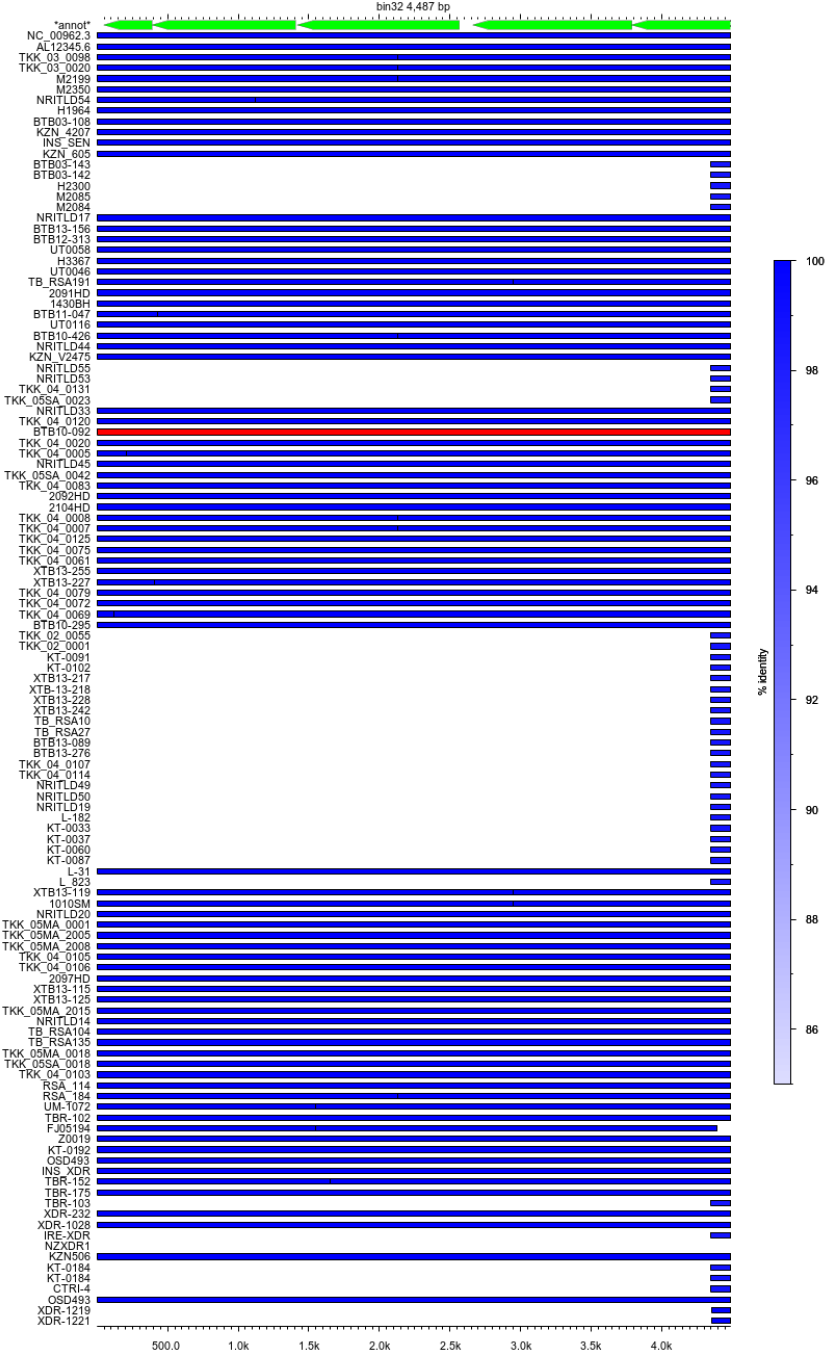
AGE graph output showing Bin 32; total size 4487 bp segment, completely deleted sequences in Beijing lineage are CRISPR-associated endo-ribonuclease cas2 (*Rv2816c*); CRISPR-associated endonuclease cas1 (*Rv2817c*); CRISPR type III-a/mtube-associated protein csm6 (*Rv2818c*); CRISPR type III-a/mtube-associated ramp protein csm5 (*Rv2819c*) and partially deleted CRISPR type III-a/mtube-associated ramp protein csm4 (61.5%) respectively

**Table 2.**
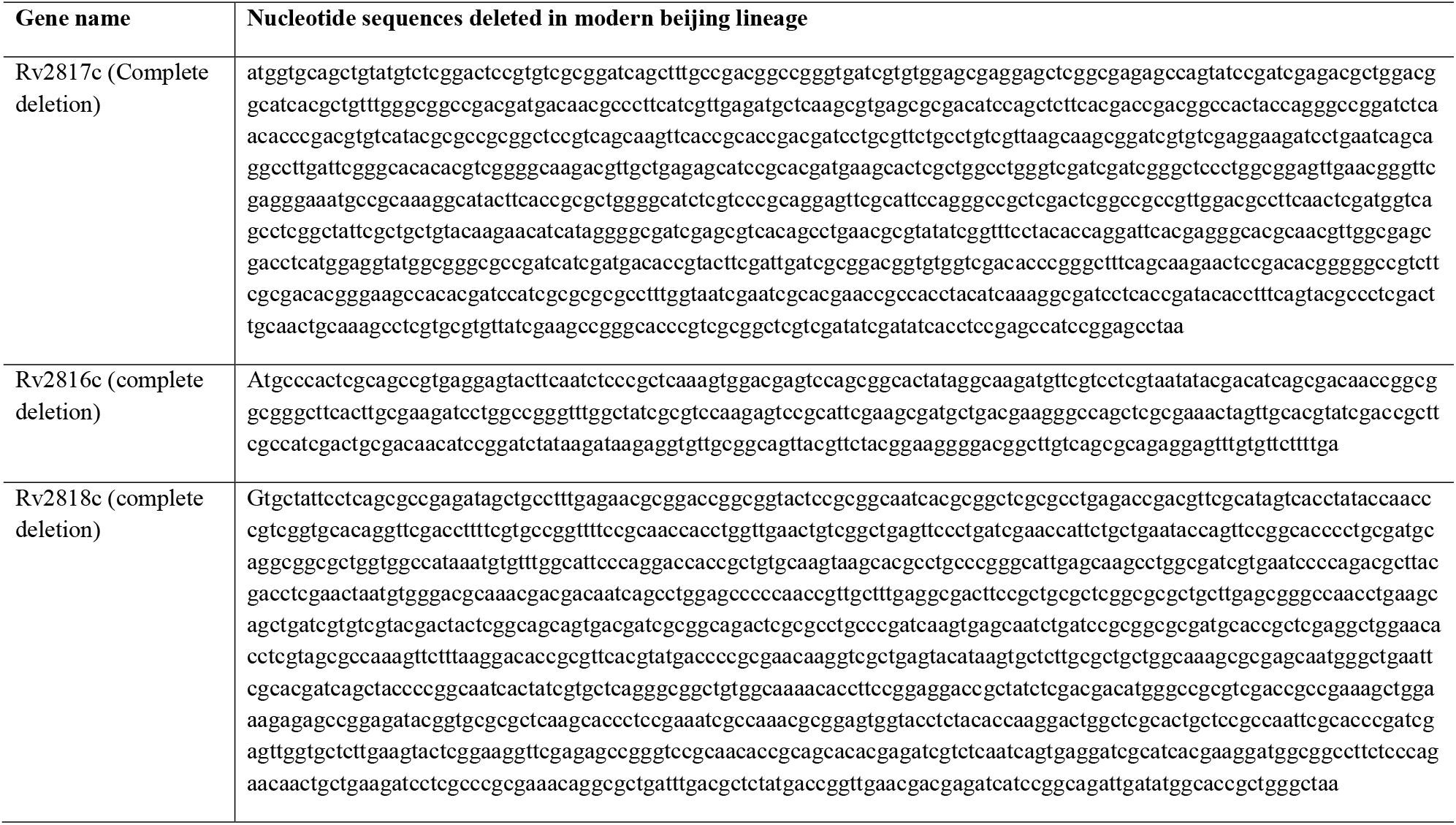
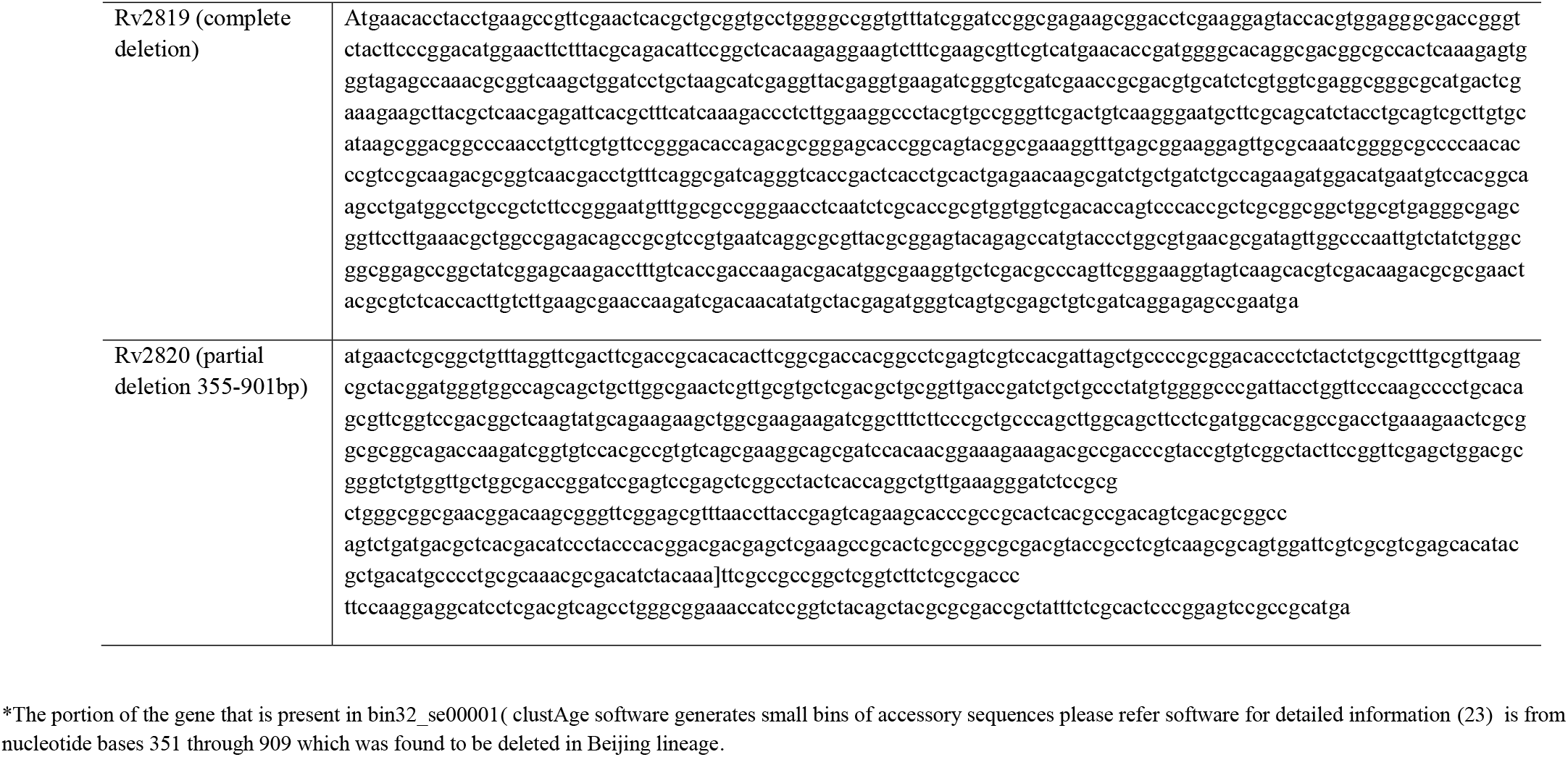
Showing Sequence elements of CRISPR-associated endo-ribonuclease cas2 (*Rv2816c);* CRISPR-associated endonuclease cas1 (*Rv2817c*); CRISPR type III-a/mtube-associated protein csm6(*Rv2818c*); CRISPR type III-a/mtube-associated protein csm5 (*Rv2819c*) and CRISPR type III-a/mtube-associated ramp protein csm4 (*Rv2820c*) deleted among modern Beijing lineage taken in assortment for pangenome analysis.

### Standardization of PCR for validation of sequences on clinical isolates

Designed primers were used for convention PCR for validation of sequences on culture isolates. Designed primers specific for CRISPR sequences are mentioned in **(Table 3)**. The PCR precisely amplified the corresponding targets from DNA isolated from the 50 non-Beijing strains and control strain H37Rv. No amplification was seen among the 50 DNA samples isolated from Beijing isolates **(Fig 9A, 9B)**. Spoligotyping patterns of isolates used for evaluation are mentioned in **(Table 4).**

**Fig 9A.**
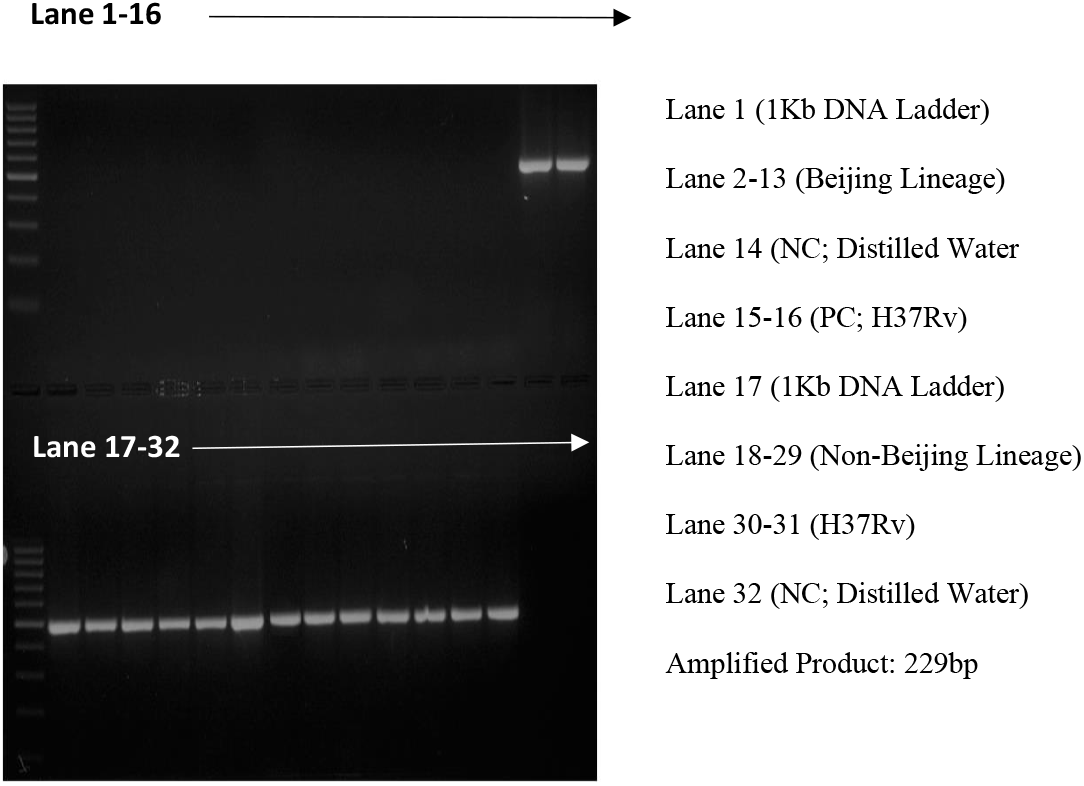
Representation pic showing CRISPR Cas2 (*Rv2816c*) sequence elements deleted in Beijing

**Fig 9B.**
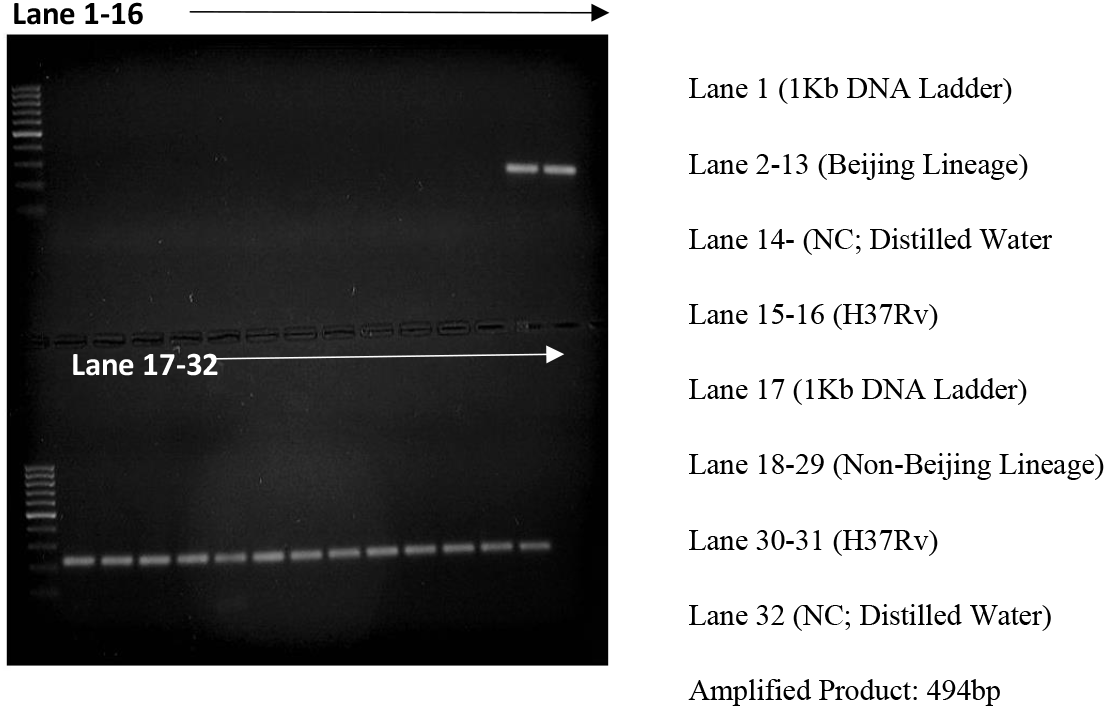
Representation pic showing CRISPR Cas1 (*Rv2817c*) sequence elements deleted in Beijing

**Table 3.**
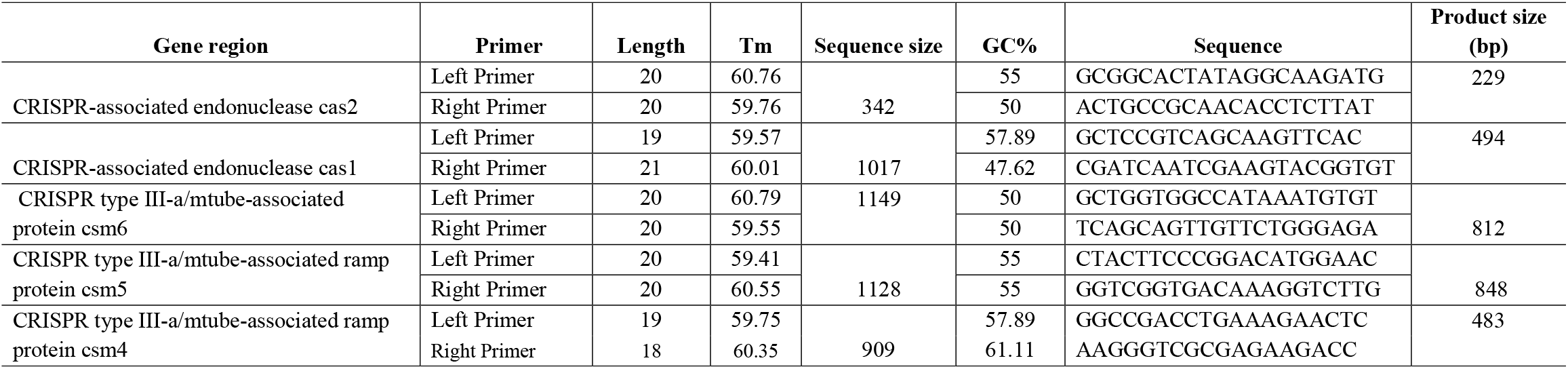
List of primers designed against CRISPR genomic elements using Primer 3 software.

**Table 4.**
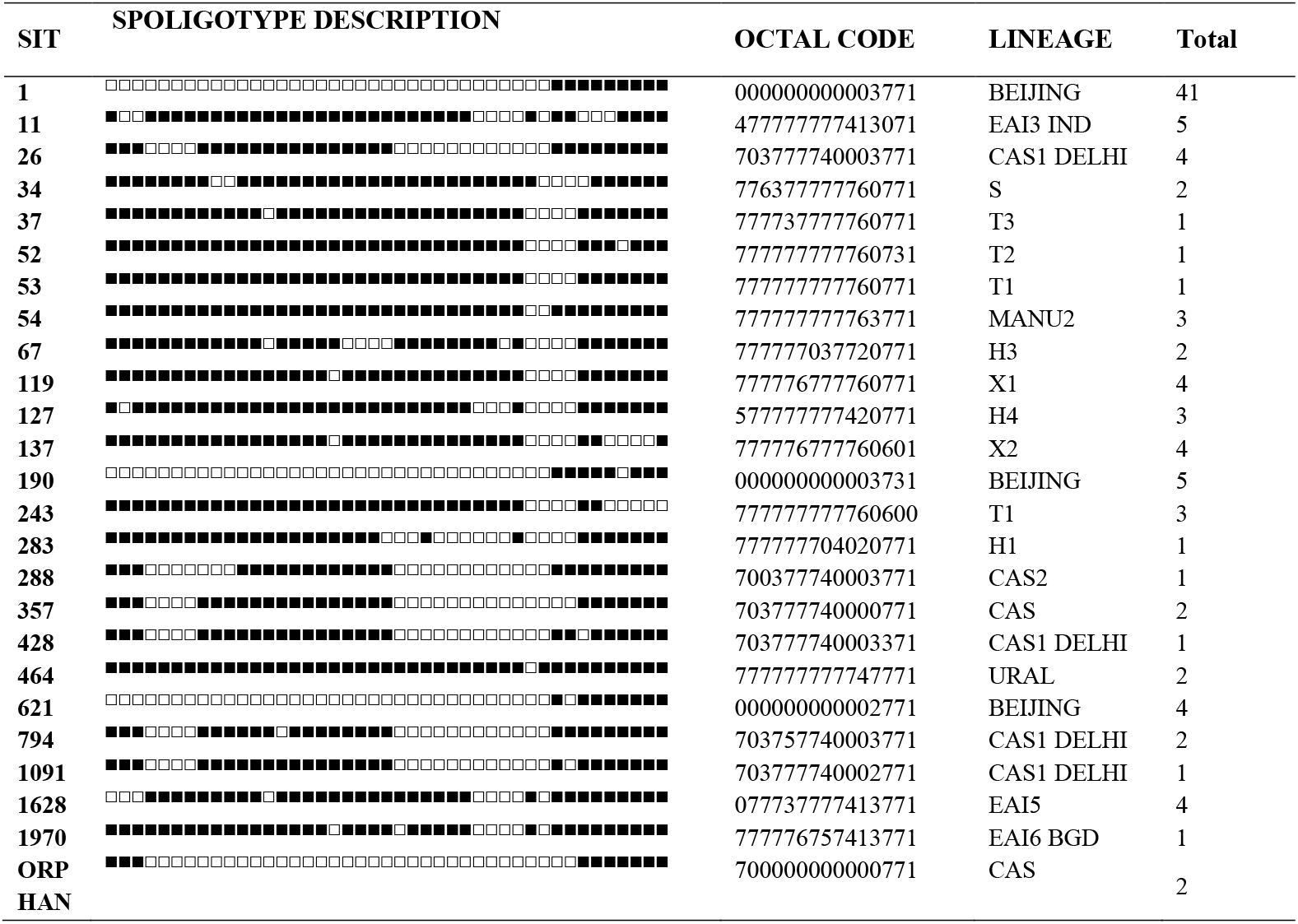
Spoligotype patterns of clinical isolates (50 Beijing and 50 Non-Beijing) used for evaluation of CRISPR sequences.

## Discussion

The main causative agent of TB in humans *M. tuberculosis sensu stricto* and of major concern is uncontrolled spread of drug resistance in developing countries (30). Total of four lineages (lineage 1-4) have been recognized within the *M. tuberculosis* species showing difference in characteristics in terms of evolutionary position, transmissibility, drug resistance, host interaction, latency, and vaccine efficacy (31). With growing evidence, it is known that genetic diversity of *M. tuberculosis* may have significant clinical consequences and sub-lineages were also reported to show similar variation characteristics especially Beijing sub-lineage (East Asian, Lineage 2). During evolution, this strain have been proposed to possess selective advantages, in contrast to other *M. tuberculois* lineages, which resulted in drug resistant TB outbreaks, increasing in population size especially in settings with distinct levels of TB incidence levels, rapid progression of disease after infection and unfavorable treatment outcome (7, 18, 19, 30–33). These findings make it crucial to understand what deviations had been altered in genome of Beijing lineage during the evolution in comparison to other lineages, resulting in excessive virulence of the strain.

To gain insights, studies have been performed to find evolutionary markers specific to Lineage2 which resulted in identification of polymorphism in noise transfer function region (NTF) locus, mutT2 and mutT4 genes, in some strains named as modern beijing sub-lineages which were predicted to be more virulent than ancient or prototype Beijing lineages (6, 34– 36). Moreover, other markers like mutation in hspX gene, deletion of *Rv0279c* in some beijing isolates gene and RD207 in all beijing isolates (*Rv2815c-Rv2820c*) were also interpreted (37, 38). As, SNPs are consistent and phylogenetically useful markers, since the low sequence distinction and lack of horizontal gene transfer in *M. tuberculosis* make independent recurrent mutations unlikely (39). We thus used comparative genomics approach to provides insight into features of shared/ unique coding sequences across different lineages of *M. tuberculosis* (23).

As expected, the predicted core genome size of *M. tuberculosis* genome repertoire was 81% of total pan-genome size for hard core genome and 95% for soft core genome representing highly conserved nature of this bacterium. In order to determine extra genes that are added in each newly sequenced genome of *M. tuberculosis* we used the concept of open/closed pan-genome (40). We predicted *M. tuberculosis* genome taken in assortment as an “open” pan-genome (as the average number of genes with each new genome shows no sign of plateauing) which specify that each new genome sequenced will provide new/novel genes, and overall increase the size of pan-genome (16). These findings correlate with previous pan-genome finding performed on Mycobacterium species (41, 42) [39]. Although, this approach of determining “open” pan-genome is mathematical extrapolation from the available sequenced genomes, however, it supports the fact that some species have tremendously flexible genetic content (40). This open/finite pan-genome implies the number of distinct genes found in *M. tuberculosis* strains is infinite as opposed to finite number of genes in a closed and thus increased emerging rate of drug resistance **(Fig 2 & Fig 3)** (41).

We found a major proportion of estimated CDS (in hard core genome) dedicated to metabolism [which consists of sequences mostly related with energy production and conversion (C), Amino acid transport and metabolism (E) and Lipid transport and metabolism (I)] **(Fig 4A & Fig 4B)**. This shows, that these sets of CDS in *M. tuberculosis* are conserved under selective pressure during its long-term interactions with its human host. The CDS associated with metabolic function may have major role in mycobacterial persistence, host pathogen struggle for nutrients, immune recognition and can be target for drug discovery [41] **(Warner, 2015)**. Average accessory genome sequences covered almost 15% of total genome repertoire, and functions of the CDS were mainly dedicated to category of poorly characterized or unknown function (S) followed by metabolism [coenzyme transport & metabolism (H)], and cellular processes & signaling [cell wall membrane envelope (M)] **(Fig 4A & Fig 4B)**. Among the poorly characterized CDS were mainly hypothetical proteins, PPE family protein, PE-PGRS family protein, PE family protein and mobile genetic elements. *M. tuberculosis* genomes containing PE/PPE family proteins have been reported as polymorphic having role in bacterial virulence and advances have been used towards the expansion of these family proteins for vaccine development (43). With such strain diversity as observed in our genome assortment taken in our study among PE/PPE family proteins there are chances for negative vaccine effectiveness however, more studies are required to prove the statement (44) (McEvoy et al., 2012).The accessory genome sequences may result in providing emergence of new functions, strain variations and understanding how it manages to survive in different niches, drug pressure which can give a reflection of its life style characteristic associated with virulence or resistance to antibiotics, it may be adapting during the course of evolution.

Unique genomic fractions were also observed in *M. tuberculosis* genomes, and were related to amino acid transport and metabolism (E), followed by Replication, recombination and repair (L) and Translation, ribosomal structure and biogenesis (J). These findings also provide information for the uptake of unique/novel genes in order to compensate the cost fitness due to antibiotic pressure. We observed majority of drug resistant *M. tuberculosis* genes were found in predicted hard-core genome except pncA (115/121 isolates). Thus, pncA gene mutations may have lower sensitivity in detection in 100% of *M. tuberculosis* strains and can be detected in a clear mainstream (>90%) of PZA-resistant strains, which has also been reported previously (45).

Our main findings during the pan-genome analysis in our collection of 121 *M. tuberculosis* genomes, we found CDS absent in modern Beijing lineage viz; CRISPR-associated endoribonuclease cas2 (100%); CRISPR-associated endonuclease cas1 (100%); CRISPR type III-a/mtube-associated protein csm6 (100%) and CRISPR type III-a/mtube-associated ramp protein csm4 (61.5%) respectively. However, we didn’t find these deletions in two genomes of Lineage 2, these two strains have been reported as prototype Beijing like harboring an ancestral-spoligotype, which is close to the Beijing clade of East Asia lineage with Spoldb4 international data base code as 246 and 643 (46). CRISPR-Cas system in *M. tuberculosis* associated with Cas1 and Cas2 genes perform endogenous DNA-repair along with a Type III A (CSM) effector arrangement, providing adaptive immunity to bacteriophages and plasmids. Deletion of the CRISPR-Cas system with associated Cas1 and Cas2 genes along with Type III A (CSM) among Beijing lineage strains could suggest more defective DNA-repair genes in such strains resulting in additional variability (47). This could predispose the lineage to development of drug resistance and transmission in the community. Such sequence markers could be useful in geographical regions where predominance of Beijing lineage is suspected. Beijing lineage that is vulnerable to first- and second-line TB drugs and has role in MDR-TB transmission. We also validated the CRISPR sequences that we predicted to be deleted in modern beijing lineages on lab isolates having different spoligotype patterns **(Table 4)** using conventional PCR method, which resulted in 100% sensitivity and specificity. This will facilitate molecular epidemiological studies and may contribute in the identification of virulent Beijing strains. Additional evidences including expression of cas1 and cas2 gene across *M. tuberculosis* lineages need to be verified to conclude that deletion of these genes lead to increased sensitivity to DNA damage resulting it in a potential phenotype mutator.

## Conclusion

We conclude *M. tuberculosis* with an open pan-genome, presence of unique genome sequence fractions which may have significant role to persist in host, tolerating antibiotic pressure and developing drug resistance. Moreover, we found modern Beijing strains to have accessory sequences elements deleted which may have role in virulence and adaptation among these strains. Further, in-depth gene expression studies are required to understand the role of such sequences in Beijing Lineage.

## Acknowledgement

A fellowship to S.B.R. from the All India Institute of Medical Sciences, New Delhi, India (reference number P-2012/12452) is likewise acknowledged. This study was supported by a grant from the Department of Biotechnology of the Government of India (BT/538/NE/TBP/2013) and the Indian Council of Medical Research (5/8/5/41/2016/ECD-I).

